# Consensus transcriptional states describe human mononuclear phagocyte diversity in the lung across health and disease

**DOI:** 10.1101/2020.08.06.240424

**Authors:** Joshua M. Peters, Paul C. Blainey, Bryan D. Bryson

## Abstract

Monocytes, dendritic cells, and macrophages, commonly referred to as mononuclear phagocytes (MNPs), are innate immune cells capable of adopting diverse homeostatic and pathogenic phenotypes. Recent single-cell RNA-sequencing studies across many diseases in the lung have profiled this diversity transcriptionally, defining new cellular states and their association with disease. Despite these massive cellular profiling efforts, many studies have focused on defining myeloid dysfunction in specific diseases without identifying common pan-disease trends in the mononuclear phagocyte compartment within the lung. To address these gaps in our knowledge, we collate, process, and analyze 561,390 cellular transcriptomes from 12 studies of the human lung across multiple human diseases. We develop a computational framework to identify and compare dominant gene markers and gene expression programs and characterize MNP diversity in the lung, proposing a conserved dictionary of gene sets. Utilizing this reference, we efficiently identify disease-associated and rare MNP populations across multiple diseases and cohorts. Furthermore, we demonstrate the utility of this dictionary in characterizing a recently published dataset of bronchoalveolar lavage cells from COVID-19 patients and healthy controls which further reveal novel transcriptional shifts directly relatable to other diseases in the lung. These results underline conserved MNP transcriptional programs in lung disease, provide an immediate reference for characterizing the landscape of lung MNPs and establish a roadmap to dissecting MNP transcriptional complexity across tissues.

## INTRODUCTION

Monocytes, dendritic cells, and macrophages represent a group of innate immune cells commonly referred to as the mononuclear phagocyte system (MPS) (Guilliams et al., 2014; Jenkins and Hume, 2014). These cells are derived from either fetal, yolk sac, or adult hematopoietic progenitor cells. These cells are responsible for a range of critical functions, such as phagocytosis of foreign bodies, presentation of antigens, cytokine secretion, and tissue maintenance. (Blériot et al., 2020; Nutt and Chopin, 2020). To perform these functions, these cell types execute diverse and distinct transcriptional programs in response to local and systemic cues (Jaitin et al., 2019; Maier et al., 2020; Okabe and Medzhitov, 2016; Reyes et al., 2020). This transcriptional diversity has been particularly appreciated in macrophages, where the classically-defined dichotomous spectrum has been progressively expanded in favor of a complex spectrum both *in vitro* and *in vivo* (Gautier et al., 2012; Li et al., 2019; Rajab et al., 2019; Sica and Mantovani, 2012; Xue et al., 2014). Defining the landscape of transcriptional states occupied by these cell types will be critical to uncovering, interrogating, and targeting the molecular signals and factors responsible for driving these states (Chow et al., 2011; Ginhoux et al., 2016; Hashimoto et al., 2011).

The diversification of macrophages across tissue niches has been defined by specific transcription factors, chromatin architecture, and gene expression programs (Blériot et al., 2020; Lavin et al., 2014). However, the diversification within tissue niches has been a target of more recent work, focusing on four main driving signals: origin, the local microenvironment and its immune state, and time (Blériot et al., 2020). The lung is one such tissue niche containing a dynamic MNP population, including various resident macrophage populations, in addition to monocytes, recruited macrophages, and a host of dendritic cell subsets. Local macrophage populations can be replenished by blood monocytes in mice, which can progressively differentiate and acquire macrophage lineage and lung transcriptional markers (Liu et al., 2019; Misharin et al., 2017). The remodeling of the lung myeloid compartment as a whole is a recurring theme across lung diseases based on conventional markers (Baharom et al., 2017). Most recently, large shifts in MNP populations have been described in COVID-19 (Liao et al., 2020). However, the conserved and distinct aspects of this remodeling has not been fully explored.

Single-cell RNA-sequencing (scRNA-seq) has been used to profile millions of cells across human tissues and diseases, providing the opportunity to better define the complex phenotypes MNPs can occupy across health and disease (Svensson et al., 2019). Broadly, numerous studies have identified unique monocyte, dendritic cell, and macrophage subsets associated with various tissue and sample classifications (Jaitin et al., 2019; Maier et al., 2020; Masuda et al., 2019; Ramachandran et al., 2019; Reyes et al., 2020; Smillie et al., 2019). The lung has been the subject of extensive profiling across multiple studies and diseases (Habermann et al., 2019; Lambrechts et al., 2018; Laughney et al., 2020; Madissoon et al., 2019; Mayr et al., 2020; Morse et al., 2019; Raredon et al., 2019; Reyfman et al., 2019; Travaglini et al., 2019; Vieira Braga et al., 2019; Zilionis et al., 2019). These studies have revealed substantial diversity of the MNP compartment, including SPP1+ macrophages (Morse et al., 2019; Reyfman et al., 2019), activated dendritic cells (Laughney et al., 2020), and chemokine-expressing macrophages (Zilionis et al., 2019). Moreover, two recent studies compared MNP populations between humans and mice in great detail, showing broad correspondence between states but caution when identifying more specific states and genes (Leach et al., 2020; Zilionis et al., 2019). While these studies have reinforced intra-niche diversity, the similarities and differences of the niche between diseases and cohorts remains unexplored.

To address the aforementioned gaps in our knowledge, here we harmonize consensus transcriptional signatures across 12 studies spanning 149 donors in order to advance our nomenclature and understanding of myeloid diversity. Utilizing two complementary methods, we identify conserved gene sets describing discrete cell states and more continuous gene programs, observing similar MNP populations in multiple cohorts and diseases. We capture both established MNP populations, such as blood monocytes, tissue-resident macrophages, and plasmacytoid dendritic cells, while also providing additional support of emerging signatures, such as intermediate, fibrotic, and chemokine-expressing macrophages states. We use this reference to dissect disease-associated trends in cancer, interstitial lung disease, and viral infection, highlighting efficient interrogation, interpretation, and support of lung MNP scRNA-seq profiles in the context of many diverse studies. This reference can be utilized similarly to common enrichment databases, but offers transparent sources and focuses on consistent, highly-detected genes in single-cell technologies. Our results establish a high-dimensional transcriptional reference for MNPs revealing the drivers of transcriptional heterogeneity in the lung and enable rapid contextual description of published and new data.

## RESULTS

### Standardized processing of lung MNPs generates robust markers and variable genes

To analyze the lung MNP diversity, we obtained, processed, and annotated cellular profiles across 12 studies and 149 donors split across 160 technical batches. These samples profile idiopathic pulmonary fibrosis, non-small cell lung cancer, and systemic sclerosis-associated interstitial lung disease, among others (Table S1 and S2). We only included droplet-based single-cell RNA-sequencing technologies. To maintain the integrity of disease-specific effects within each dataset, we split the count matrices into healthy and disease count matrices, creating 18 total datasets representing the expression per gene per cell for each cohort of samples. Count matrices were processed using a semi-automated standardized pipeline, applied consistently to all datasets to ensure uniform computational transformations (Figure 1A, S1A-S1C, **Methods**) (Germain et al., 2020; Hie et al., 2020; Stuart et al., 2019). First, we identified high-quality cells by filtering on technical metrics such as library size and percentage of mitochondrial reads. Significant batch effects were observed in all datasets, so we employed Harmony to correct for these effects in the PCA-generated low-dimensional space, which was used for downstream clustering and visualization only (Korsunsky et al., 2019; Traag et al., 2019). We found that the proportion of variance explained by principal components was similar across datasets (Figure S1D), which may be expected due to the similar cellular populations. After this initial pre-processing, we sought to broadly annotate cell populations and identify monocytes, dendritic cells, and macrophages.

**Figure 1.**
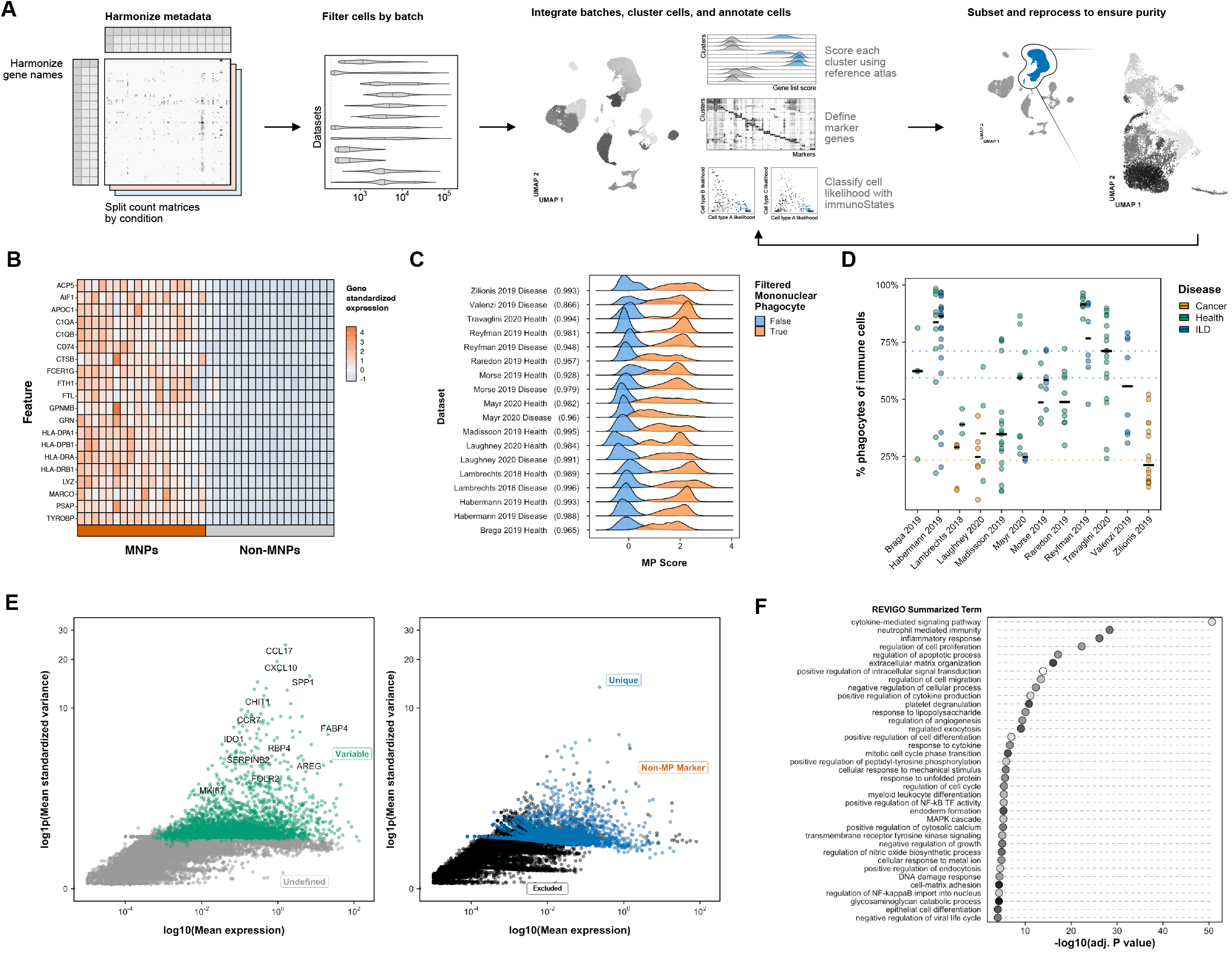
Collation and identification of mononuclear phagocyte single cell transcriptomic profiles reveals robust markers and variable genes. **(A)** Data are harmonized, filtered, batch-corrected and annotated using a standardized pipeline MNP profiles are identified using reference gene markers and classifiers. **(B)** 20-gene signature of the most predictive genes for MNP clusters across datasets. Marker genes are identified by testing each MNP cluster against all non-MNP cells within that dataset. Each column represents a dataset, labeled as either MNPs or non-MNP cells. **(C)** MNP signature score (average expression of genes above baseline expression) is highly predictive across datasets between MNP and non-MNP cells. AUCs range from 0.866 (Valenzi 2019 Disease) to 0.996 (Lambrechts 2018 Disease). **(D)** Phagocyte proportion varies widely within and between datasets and diseases (summarized as interstitial lung disease (ILD), healthy, and cancer). Black bars represent median percentage. **(E)** Average variance (log(standardized variance + 1)) and average expression (log(mean expression)) of all genes within the mononuclear phagocyte compartment across all datasets highlight consistent highly variable genes. Consensus variable genes, colored green, represent the top 3000 well-detected, consistently genes ranked by the number of datasets the gene was independently identified as variable. Selected high variance genes are labeled. Genes colored blue represent unique variable genes not identified in other datasets. Orange genes represent variable genes identified in the MP population that are strong predictive markers of non-MP cell types. Excluded genes represent a combination of unannotated and receptor genes. **(F)** Variable genes are enriched (GO Biological Process, enrichR) for diverse functions, semantically summarized using REViGO (similarity allowed 0.7).

We used multiple, complementary procedures to annotate clusters, including inspecting differentially-expressed genes, classifying cells using the immunoStates expression reference (Vallania et al., 2018), and scoring cells based on a recent healthy lung atlas (Travaglini et al., 2019). To score cells, we extracted the top 20 defining genes (e.g. SFTPC for alveolar epithelial cells, COL1A1 for lipofibroblasts, MS4A1 for B cells, and S100A8 for classical monocytes) from each major cell type defined in this atlas. These scores represent the expression of the gene set above baseline expression (Figure S1E) (Tirosh et al., 2016). To annotate clusters in an unsupervised manner, we utilized a voting procedure by assigning each cell a class label (e.g. Immune), subclass (Lymphocyte), and cell type label (e.g. B cell) based on the maximum score. Clusters were broadly annotated based on which assigned cell types were most frequent (Figure S1F), providing both a cluster annotation and a measure for the strength of that annotation.

To specifically identify clusters representing MNPs, we calculated a total MNP score for each cell, representing the difference between all MNP and all non-MNP scores. We defined a threshold score from this bimodal distribution by fitting two Gaussian distributions to these scores (**Methods**). Using this threshold, we extracted high-confidence clusters based on a median MNP score above threshold and a low frequency of non-MNP cell labels from the voting procedure. These clusters were processed and filtered in a semi-supervised manner to ensure high-quality MNP profiles based on all three metrics described here (**Methods**). Retained MNP clusters consist of cells classified as MNPs by Bayesian classifiers, show high and broad expression of previously published lung MNP markers, and are not defined by technical or non-MNP gene markers. After filtering, these cells were again reprocessed. This procedure produced concordant annotations with available, published annotations (Figure S1G). Overall, following gating and quality control filters, we analyzed 180,629 MNP scRNA-seq profiles, out of 561,390 total profiles, from 160 batches, 149 donors, and 12 studies (Figure S1H).

We next sought to define a signature of MNPs built using single-cell RNA sequencing data that was robust across multiple datasets. By determining the most predictive genes for each MNP cluster versus all non-MNP cells, we defined a 20-gene signature that performed well in broadly classifying MNPs (AUC range 0.87-0.996 based on MNP annotation described previously) (Figure 1B and 1C). This signature included well-known mononuclear phagocyte markers, such as the MHC Class II molecules (HLA-DRA, HLA-DRB1, CD74) and recently identified markers from purified populations (TYROBP, FCER1G) (Dang et al., 2020), as well as genes associated with other various functions, such as iron regulation (FTH1, FTL), complement (C1QA, C1QB), and lysosomal processing (CTSS, CTSB, LYZ, PSAP). ACP5 was also identified, which encodes a critical regulator of osteopontin (SPP1), which has been implicated in lung cancer and fibrosis (Figure 1C) (Kim et al., 2020; Reyfman et al., 2019). MNPs represented a large proportion of leukocytes in each technical batch, but varied widely across samples (Figure 1D).

We next sought to broadly delineate the dominant highly variable genes underpinning diversity within these cell populations. Single-cell analyses typically leverage this subset of genes to focus on the most differentiating signals and reduce noise from uninformative genes. Consistent, variable genes represent critical molecular features responsible for, or a consequence of, intra-tissue cellular diversity. We identified a set of 3,665 consensus highly variable genes that underpin the heterogeneity in the lung MNP compartment (Figure 1E and Table S3). These genes included well-established immunological markers and targets, such as IDO1, CXCL10, CCR7, and FOLR2 as well as less canonical genes FABP4, CHIT1, RBP4, SERPINB2, SPP1, SEPP1, and AREG. These genes should be studied further for their role in shaping MNP population structure. Robust, highly variable features within the MNP compartment in the lung could be a consequence of the underlying diversity or have a role in creating that diversity. Gene set enrichment of these consensus variable genes using GO Biological Process database revealed a diversity of functional associations related to the host of roles MNPs perform, most notably in angiogenesis, metabolic homeostasis, and mechanical extracellular interactions (Okabe and Medzhitov, 2016). Indeed, the enrichment of a diverse functional set suggests that the transcriptional diversity is indicative of the many functions MNPs perform, supportive of the “variation is function” hypothesis (Dueck et al., 2016).

### Pan-cluster comparison reveals conserved cell subsets and markers

Previous single-cell studies identified distinct transcriptional subpopulations in the lung MNP compartment, including macrophage, monocyte, and dendritic cell subtypes. To date, our understanding of the similarities across these subpopulations and studies remains limited. We first investigated whether distinct MNP transcriptional states were conserved and defined similarly across studies. To systematically compare cell states across each dataset, we established clusters, representing cells with similar expression profiles, and their respective gene markers within each dataset. Choosing the resolution of clustering remains an open and consistent challenge in single-cell RNA-sequencing analysis, so we focused on clustering cells at higher resolutions (creating more clusters with fewer cells) and retaining only clusters strongly described by multiple, highly-expressed genes based on various metrics such as the log fold change (**Methods**). This enabled us to capture the most distinctive, well-described cell states in each dataset (n = 295 total clusters, n = 267 with strong markers) described by up to 50 unique gene markers (Table S4). For example, in two studies, Lambrechts et al. and Laughney et al., clusters emerged that were defined by strong inflammatory markers, such as CXCL10 and GBP1.

To determine the extent to which the diversity of the MNP populations are conserved across samples and diseases, we calculated the similarity of all states using the Jaccard distance, which represents the overlap of two gene lists (Jaccard distance = 0, perfect overlap) (Kinker et al., 2019). We focused on 151 clusters that reached a minimum Jaccard index of 0.2 (Jaccard distance of 0.8) with at least one other cluster (**Methods**). These states were then hierarchically clustered to identify partitions of similar cell states defined by similar gene markers across various studies (Figure 2B). To determine the number of partitions, we defined a cutoff based on cluster distances and agreement with other methods, resulting in 21 partitions (Dynamic Tree Cutting, adjusted Rand index = 0.92 where 1 is perfect agreement; Cluster Bootstrapping, ARI = 0.94, Figure S2A-S2C) (Hennig, 2007; Langfelder et al., 2008). These partitions contained clusters from multiple datasets (Shannon diversity between 1.5 and 2.5, Figure 2C). Partitions with low diversity and reproducibility were filtered out and removed from downstream analyses (partitions 6, 14, 19, 21). To further confirm the similarity of cell states within partitions, we utilized *scmap*, an unsupervised, high-performing classifier, to calculate a similarity metric for all pairwise combinations of all clusters (Kiselev et al., 2018). Consensus partitions were significantly more similar than random equally-sized partitions (P < 0.001, Figure S2D, **Methods**). We define these distinct partitions as consensus markers denoted as “M[X]” where X is an arbitrarily assigned number (Figure S2E, Table S5).

**Figure 2.**
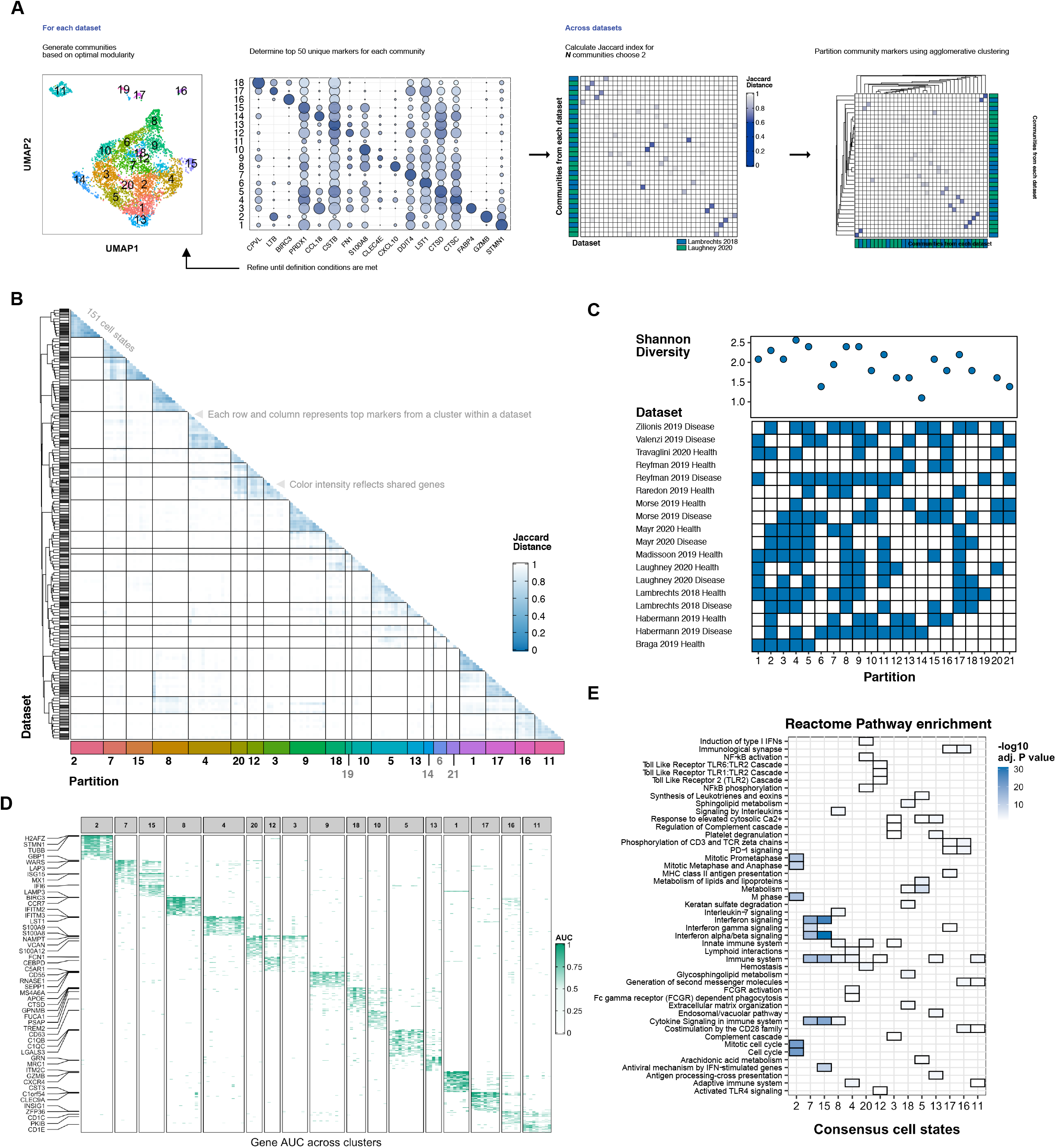
Cluster-defining genes are similar across datasets and represent conserved cell subsets. **(A)** Clusters are defined and refined based on modularity and marker characteristics. Markers for each cluster are compiled and the Jaccard distance is calculated between all intra- and inter-dataset comparisons of clusters.Example similarity analysis utilizes disease samples from Lambrechts and Laughney studies. **(B)** Hierarchical clustering of the Jaccard distance matrix between all clusters’ marker genes reveals distinct partitions of similar cell states. Markers are defined by the Wilcoxon rank sum test, auROC, and a minimum second-to-max log fold change of 1.05. Matrix is clustered based on agglomerative hierarchical clustering (k = 21). Clusters with less than 5 markers or no matches above a Jaccard index of 0.2 are not included. **(C)** Each partition contains cell states from different studies. Each filled tile represents the presence of that dataset within that partition. Shannon diversity is plotted above, highlighting values primarily between 1.5 and 2.5. **(D)** Partitions are defined by high, median auROC values. The top 3 marker genes defined by clusters identified in (A) are labeled. Low quality partitions are removed. **(E)** Enrichment of programs using the Reactome Pathway Database underlines distinct cell states across MNPs. All terms are enriched at adjusted P value < 0.05. Color represents −log10(adjusted P value).

We sought to define a conserved set of marker genes describing these partitions. To define these gene markers, we utilized the auROC metric which provides a relative indication of how well expression of specific genes signifies a specific cell state as defined by cluster membership (auROC = 1, perfect classifier, Figure 2D). This revealed highly concordant marker genes within partitions, both confirming the consensus cell states as well as highlighting dominant signatures defining heterogeneity in the MNP compartment. As expected, we found prominent signatures of more familiar signatures, such as replicating cells (H2AFZ+, M2), CD14+ monocytes (S100A9+, M20), CD1C+ dendritic cells (PKIB+, M11), and plasmacytoid dendritic cells (GZMB+, M1) (Travaglini et al., 2019). These integrated signatures offer a highly detected and consistent reference. Moreover, these consensus cell states reveal multiple macrophage states. These states reflect mature alveolar-like states PPARG+ (M13) and C1QB+ (M5), as well non-tissue resident states SPP1+ (M10), APOE+ (M18) and SEPP1+ (M9) states, and immune-associated states GBP1+ (M7) and MX1+ (M15) (Reyfman et al., 2019; Zilionis et al., 2019). Additionally, this approach uncovered a rare recently described anti-tumor-associated cross-presenting dendritic cell subset (LAMP3+, BIRC3+, CCR7+), identified in both healthy and interstitial lung disease samples as well (Maier et al., 2020).

Enrichment of these gene sets using the Reactome Pathway database highlights a wealth of distinct functional processes (Figure 2E), owing to the diversity in cellular states identified. M7 and M15 are similarly associated with interferon signaling, although M7 is enriched for IFNγ signaling versus antiviral signaling in M15. M18, comprising apolipoprotein and lipid-associated genes, is enriched for sphingolipid processes suggesting a role for these molecules in population diversity. As expected, dendritic cell-like states were enriched for MHC Class II presentation and T-cell signaling (M11, M16, M17). Projection of these consensus cell states and their underlying populations using UMAP visualizes and reinforces this transcriptional state landscape (Figure S2F) seen in the MNP compartment. Indeed, projection of the auROC values for each contributing state and the resulting consensus states reveals highly distinct and coherent groups (Figure S2G). Overall, these consensus cell states, defined by robust, consistent genes, broadly define the diversity of the MNP population in a large cohort of lung samples.

### Integrated gene expression programs show similar concordance across datasets

Given the transcriptional spectrum seen in the mononuclear phagocyte compartment (Figure S3A), we utilized an orthogonal approach to capture more continuous gene expression programs (GEPs). These GEPs aim to capture coregulated genes that describe cellular behavior both within and across cell types, such as a type I interferon response or general response to stress. Non-negative matrix factorization (NMF) has been established in single-cell RNA-sequencing analysis to identify gene expression programs descriptive of cellular identity and activity (Kinker et al., 2019; Kotliar et al., 2019; Welch et al., 2019). Thus, by comparing GEPs across datasets, we aimed to identify consensus GEPs which consistently describe variation in the MNP compartment. We combined two previous approaches, consensus NMF and integrative NMF, to define 20 consensus, integrated factors representing GEPs (Figure 3A). In this approach, we perform integrative NMF across batches many times to generate robust batch-agnostic GEPs and utilize the top 5% contributing genes to each GEP for comparisons (**Methods**) (Kotliar et al., 2019; Welch et al., 2019).

**Figure 3.**
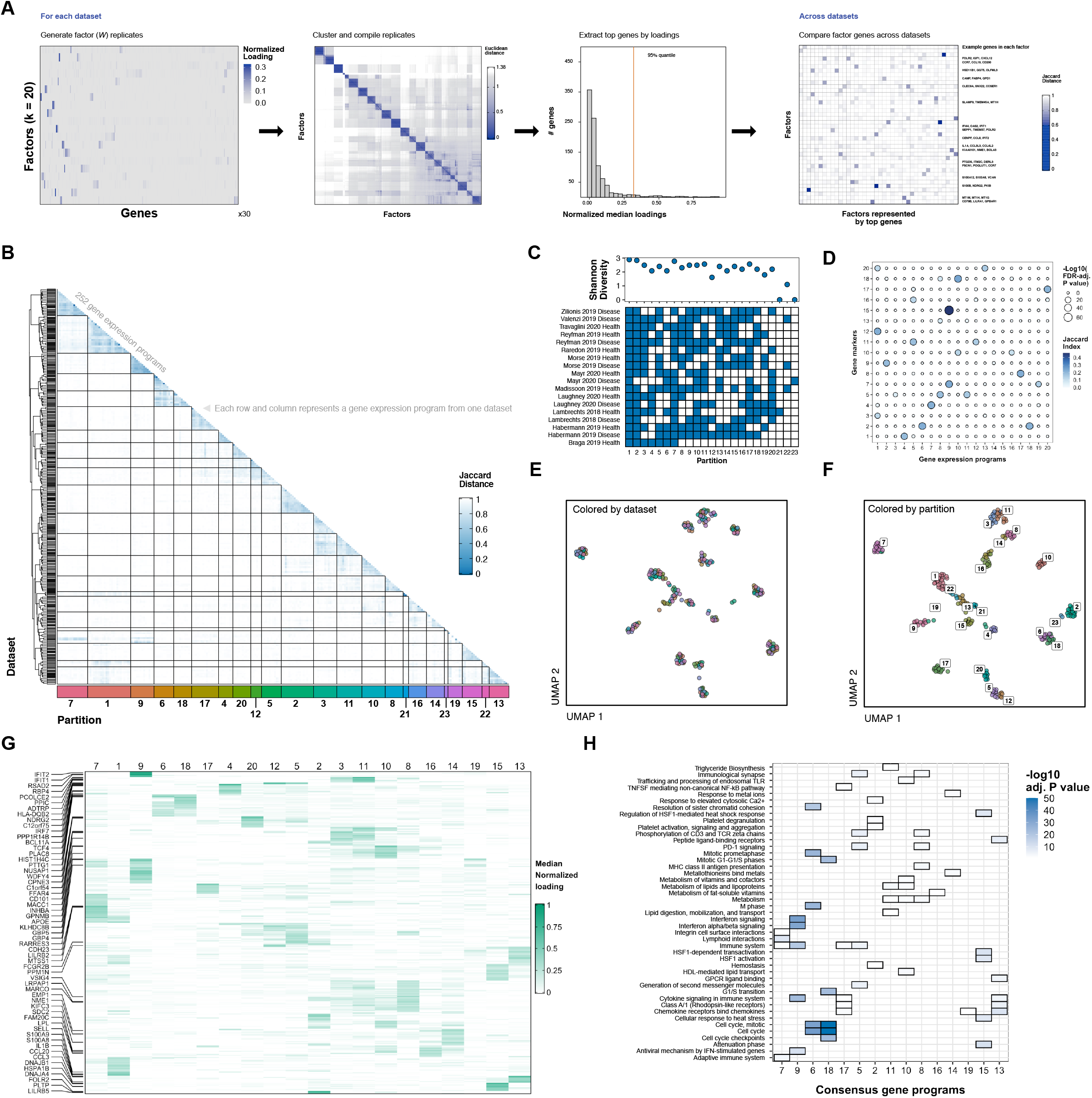
Gene expression programs are conserved across datasets, complementary to differentially-expressed genes, and describe the continuous identity and activity of phagocytes. **(A)** GEPs are identified through consensus results from iNMF. iNMF generates factors which are compiled over 30 replicates. The top 5% of genes for each compiled factor are then used for comparison using the Jaccard distance. **(B)** Factors are partitioned (k = 23) using agglomerative hierarchical clustering on all GEPs defined using procedure in (A). **(C)** Diversity indices and binary presence matrix highlights diverse dataset membership. **(D)** Comparison of consensus gene expression programs defined in this figure with consensus gene markers defined in Figure 2 underlines similar and unique gene sets. Size indicates significance of gene overlap and color represents the Jaccard index. **(E)** GEPs can be visualized in a UMAP embedding, highlighting no dataset-specific effects. Points are colored by dataset. **(F)** GEPs grouped by partition are localized in UMAP embedding. **(G)** Partitions are defined by median gene loadings. Each value represents the median loading of a gene calculated from each dataset-specific factor within that partition. **(H)** Consensus GEPs are enriched using the Reactome Pathway database. All terms are enriched at an adjusted P value < 0.05. Color represents −log10(adjusted P value).

Using the same approach described for markers, we identify 23 initial partitions, resulting in 20 partitions after inspection and filtering (Dynamic Tree Cutting, ARI = 0.97, Cluster Bootstrapping, ARI = 0.88, Figure 3B, Figure S3B-S3D). Similarly, passing partitions are highly diverse, comprising program gene sets from many datasets (Figure 3C). To define a set of genes for each partition, we selected the top 50 genes with the highest median contribution to the GEPs within the partition (**Methods**). We define these consensus program gene sets as “P[*X*]” where *X* is an arbitrary assigned number (Table S6). We compared these consensus programs to the previously defined consensus markers, revealing unique concordance between pairs of gene sets, such as P4-M1 and P2-M9 (Figure 3D). Some gene sets also show redundant overlapping, reflecting the similarity seen between certain gene sets, such as the monocyte-like marker sets defined by P1 and M3/12/20. Interestingly, P13 shows distinct similarity with M20, representing inflammasome-related genes IL1B and NLRP3. While P9 is similar to M7 and M15, P19 shows strong similarity with just M7, due to CXCL10, CXCL11, and GBPs. These gene sets provide corresponding descriptions of dominant cellular states and their underlying gene programs in the lung MNP compartment in health and disease.

Using the loadings of each gene for each GEP which represent the genes’ importance in that GEP, we visualized the programs from each dataset, producing an embedding with no discernable batch effects (Figure 3D and 3F). Programs that clustered together based on overlap of the most-defining genes also clustered distinctly within the embedding, supporting the consensus partitions. Moreover, these consensus GEPs are distinctly defined by the median gene contributions (Figure 3G), highlighting both the congruence of dominating genes in each GEP. This supports the ability to represent and analyze cellular variation from disparate datasets as GEPs derived from NMF (Welch et al., 2019).

Inspection and enrichment of the consensus programs produces similar pathways to the consensus markers, but also includes enrichment of lipid-associated processes in P10 (APOC1, APOC2, LIPA, and TREM2) and P11 (FABP3, FABP4, FFAR4, LPL, SCD). Moreover, we see the appearance of P15 associated with the heat shock protein response and P14 associated with the metal ion response. P13 is enriched for chemokine expression and GPCR signaling, described by a host of immunoregulatory genes including CCL3, IL8, IL1B, IL6, NFKB1, and TNFAIP3. Comparison to conventional M1 and M2 signatures suggests moderate correlation between signature scores (M1 with P9, P13, P17, and P19 and M2 with P2, P8, and P10) (Figure S3F). This GEP-focused approach produced similar and additional gene sets compared to the marker-based approach, providing both support for these findings and expanding our definition of underlying molecular signatures. Overall, we present two concordant lists of marker and program gene sets which describe conserved, consistent transcriptional signatures in the lung mononuclear phagocyte compartment.

### Consensus programs and markers describes disease- and cell type-associated variability

Dysregulation of MNPs has been hypothesized or shown to contribute to lung diseases included in this work (Reyfman et al., 2019; Zilionis et al., 2019), so we reasoned that we could leverage our consensus gene programs and markers to define the transcriptional programs associated with these cells during disease. We focused on comparing healthy and disease samples in a subset of 6 studies where this comparison was possible. We utilized a scoring approach that calculates a normalized expression for each gene set relative to a control gene set (Smillie et al., 2019; Tirosh et al., 2016) (**Methods**). To do so, we calculated scores for each gene set (either consensus markers or programs) across all healthy- and disease-derived cells by averaging the scaled expression for each gene and corrected for random expression of 30 control genes per gene in each gene set. This procedure generates scores for each gene set and cell, representing relative expression levels of each gene set. Across clusters within each dataset, scores for both marker and program gene sets were commonly expressed in a subset of clusters (Figure S4A). We captured expression changes using a mixed linear model on the most expressing clusters (top 2, **Methods**), identifying both disease- and health-associated gene sets with variance across datasets (Figure 4A).

**Figure 4.**
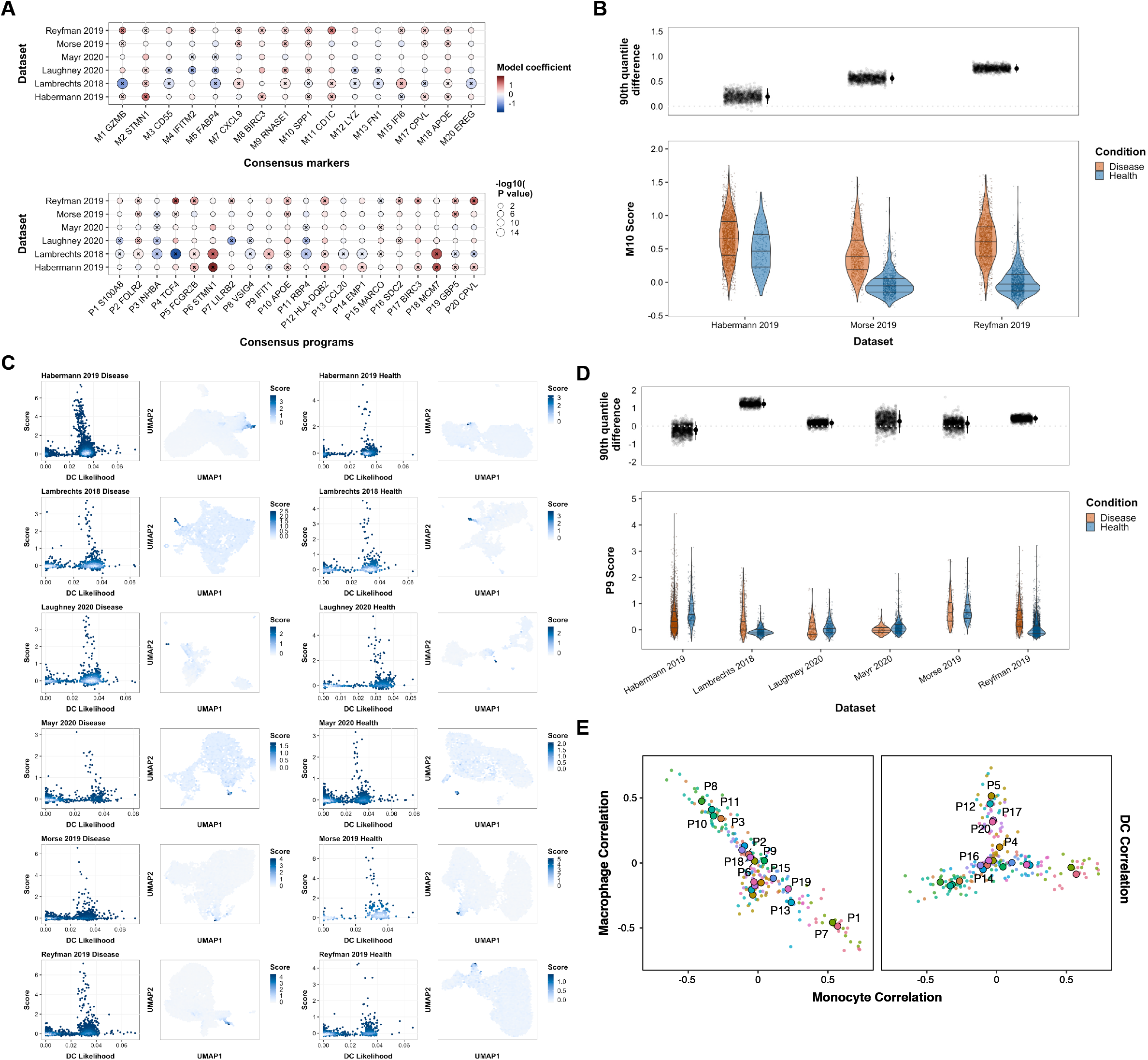
Consensus markers and programs describe disease-associated trends in the phagocyte compartment. **(A)** Mixed linear model coefficients (gene set score ~ condition + (1 | batch) for each marker and program gene set across datasets represent association with health or disease. P values are calculated using the likelihood ratio test and are Benjamini-Hochberg adjusted. Manually selected, highly associated genes are labeled for each marker and program gene set. Significant P values are marked with an “x”. **(B)** M10 scores are higher in disease samples for three datasets with pulmonary fibrosis samples. Bootstrapped 90th quantile differences between health and disease samples are also shown with median quantile intervals. Each score distribution between healthy and disease samples is significant (Mann-Whitney U test, P values < 2.2e-16). **(C)** P17 scores and DC likelihood per cell for each individual dataset identify shifts in a small subset of DCs within each dataset, representing CCR7+ activated dendritic cells. 2D hexagonal density plot of the UMAP embeddings for each data is shown right with the median score of each bin shown. **(D)** P9 scores are variably associated with disease in each dataset shown in (A). Bootstrapped 90th quantile differences also shown, similar to (B). **(E)** Pearson correlation of each program score with monocyte, macrophage, or dendritic cell likelihood underline the transcriptional spectrum describing the MNP compartment.

Using these scores, we focused on marker and program gene sets of interest. In one of the studies analyzed here, the authors identify that SPP1 expression is associated with idiopathic pulmonary fibrosis (Reyfman et al., 2019). In our analysis, we uncovered a consensus cell state M10, which includes SPP1 and is statistically associated with disease in multiple fibrosis datasets (Figure 4B). This supports their findings and reveals additional candidate genes that are robust across multiple fibrosis datasets (Reyfman, Habermann, and Morse studies) that can be collectively studied in order to define and target the dysregulated pathways responsible for this state. Additionally, we see a shift in the 90th percentile of these scores between health and disease, further suggesting an overall shift in expression (Figure 4B). P17 represents an activated, migratory dendritic cell program typically expressed in a low fraction of cells (Maier et al., 2020). In several datasets, these cells were not originally identified, so we utilized this program to identify the appearance of these rare cell types. We identified a fraction of cells in each dataset with a high P17 score and high DC likelihood based on the immunoStates reference, representing rare immunomodulatory dendritic cells (Figure 4C) (Maier et al., 2020; Vallania et al., 2018). These cells were identifiable in both healthy and disease samples. P10, which contains several genes related to the immune response including CXCL9, CXCL10, and the surface marker SIGLEC1, is associated with tumor samples from Lambrechts (Lambrechts et al., 2018). These genes were also defined by Zilionis and associated with poor prognosis (Zilionis et al., 2019). Visualization of P10 scores and a similar marker gene M7 across all 6 datasets highlights expression of these genes in a subpopulation of cells in all datasets and suggests association with disease (Figure 4D, S4B). Further focus and profiling of this population is warranted, considering the immunomodulatory mRNAs defining these cells and appearance in other studies (Giladi et al., 2020; Goldberg et al., 2018; House et al., 2019).

Given the continuous and heterogeneous expression of these programs, we sought to further annotate the association of each gene set with the likelihood of each major mononuclear phagocyte cell type. We calculated the Pearson correlation of each program score with the likelihood across all cells, revealing a ternary spectrum between monocytes, macrophages, and dendritic cells (Figure 4E). Certain programs highly associated with monocyte or macrophage likelihood did contain canonical genes associated with these cell types, such as *VCAN* and *APOE*, respectively. However, many programs showed weaker association, suggesting that many MNP populations are not well-defined by classical definitions and may represent intermediate or transitory populations (Giladi et al., 2020; Rizzo et al., 2020). Overall, we demonstrate broad utility of these gene sets to identify critical subsets, disease-associated trends, and broad population structure.

### COVID-19 myeloid dysregulation is described by MNP gene set references

We next sought to demonstrate the utility of these consensus marker and gene sets and applied our scoring methodology to interrogate phagocyte dynamics in viral infection. Given the documented dysregulation of the myeloid compartment in COVID-19, we processed recently published profiles of bronchoalveolar lavage (BAL) samples from healthy, mild, and severe COVID-19 patients (Liao et al., 2020). Viewing new profiles through this reference lens can enable quick access to biological interpretations and tools from previous work. We processed healthy, mild and severe patients separately and identified clusters using the same procedure described previously (Figure 5A).

**Figure 5.**
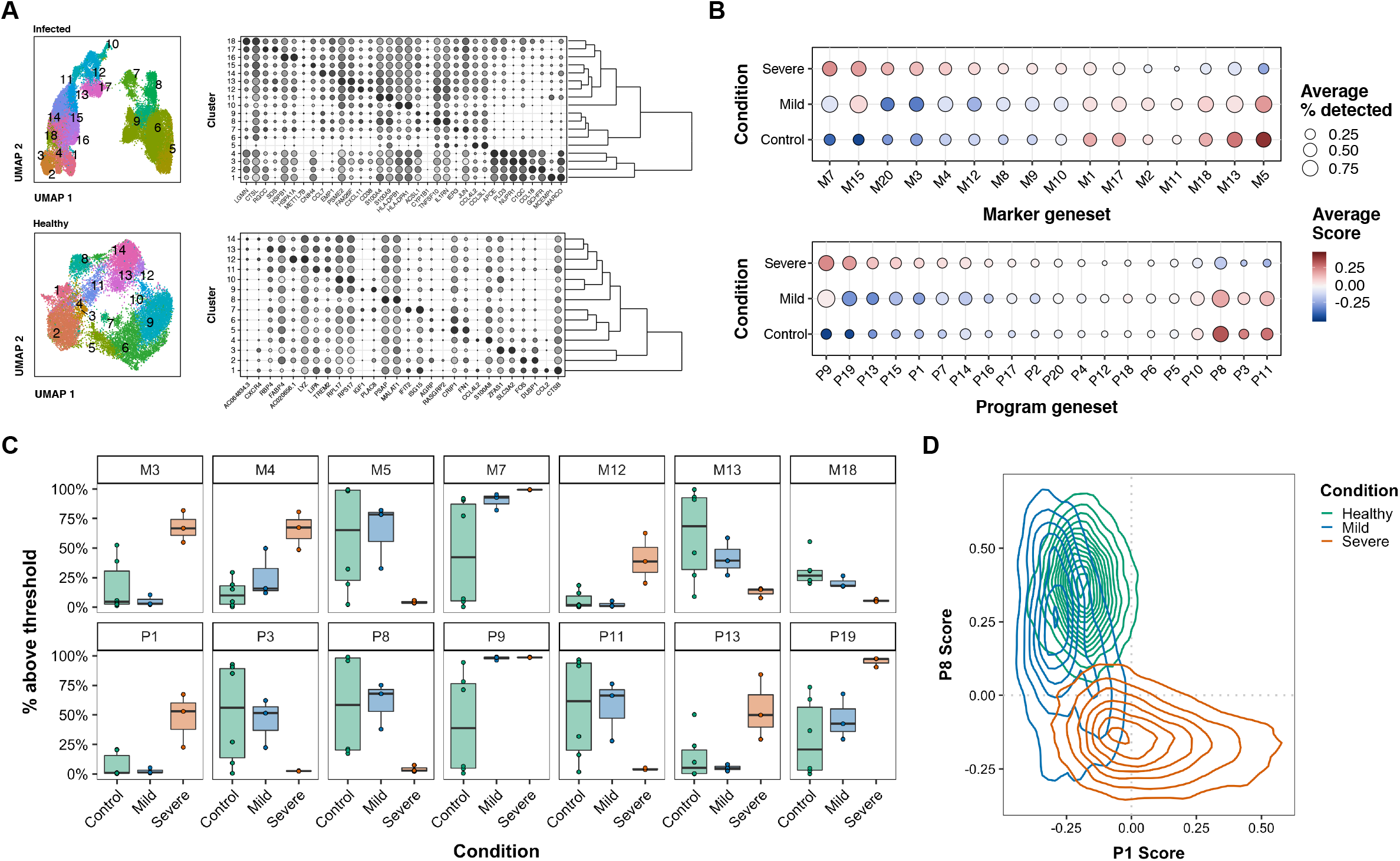
Application of consensus gene sets to COVID-19 bronchoalveolar lavage samples reveals novel aspects of phagocyte dysregulation. **(A)** Clusters, marker genes, and hierarchical trees for disease and healthy samples describe MNP populations. **(B)** Marker and program geneset scores and average detection rate across healthy, mild, and severe samples highlight high and low expression trends in infected samples vs. healthy samples. Genesets are sorted from left to right based on the average score of severe samples. **(C)** Shifts in population structure, defined by the percent of cells above threshold cells, are observed across selected marker and program scores. **(D)** P8 score (correlated with macrophage likelihood) vs. P1 score (correlated with monocyte likelihood), colored by sample condition, highlights a dominant reorganization of the lung MNP compartment.

We combined all cellular profiles and scored each cell by each consensus cell state and GEP gene set. Comparison of these scores across healthy, mild, and severe patients reveals broad increases in antiviral and inflammasome-associated GEPs identified previously in cancer, no shifts in fibrosis-related gene sets, and a decrease in GEPs describing both general and lipid-associated macrophage states (Figure 5B). M7 and P9, which similarly contain CD38, GBP1, and ISG15, and uniquely contain OAS1 and SOD2 respectively, show the highest expression in severe patients with a negative graded expression towards healthy samples. These cells expressing antiviral interferon-induced genes and lymphocyte chemokine genes have been previously annotated in cancer (Zilionis et al., 2019) and found to be associated with poor cancer prognosis. Conversely, we find that M5 (MARCO, MSR1+) and P11 (FABP3/4+), which describe an alveolar macrophage-like signature, are more prominent in healthy samples, supporting the observation of a shift away from a resident-dominated phagocyte population towards monocyte-associated populations expressing antiviral, inflammasome, and lymphotaxis-associated genes.

We next sought to describe the population dynamics within each condition using these consensus gene sets. We used an approach analogous to flow cytometry gating where we asked what proportion of cells were above a threshold. By determining the percentage of cells above an expression threshold, we quantified and visualized relative shifts in the population between each condition (**Methods**). Using this complementary approach, we find distinct infected versus healthy shifts (e.g. M7 and P9 described earlier), severe-only shifts (e.g. M3 - FCN1+, P19 - GBP5+), and graded shifts from healthy to severe (e.g. M4 - TCF4+, M13 - PPARG+, P7 - LILRB2+) (Figure 5C). Moreover, we see little expression and difference in gene sets associated with other macrophage populations (for example, M10 - SPP1+ and P2 - FOLR2+), suggesting an absence of these states. Lastly, we visualize the overall population scores of P1 (associated with monocyte likelihood) and P8 (associated with macrophage likelihood), highlighting a gradual shift of the population structure towards a monocyte-like dominated population (Figure 5D). Using this reference reveals specific identity and functional expression shifts in COVID-19 samples that can be directly contextualized in the context of broader lung populations.

### Enrichment of gene sets suggests associated molecular factors

These marker and program gene sets broadly describe phagocyte heterogeneity within the lung context. To generate candidate hypotheses for future studies which can define the underlying molecular factors for these gene sets, we performed enrichment analyses to nominate transcription factors, ligands, and receptors that may be associated with particular states.

Using the ARCHS4 transcription factor coexpression database in Enrichr, we identified several transcription factors that were significantly enriched in program gene sets versus background with prior literature support for their contribution to MNP and lung macrophage development (e.g. IRF8, MAFB, PPARG) (Figure 6A) (Lavin et al., 2014; Mass et al., 2016). PPARG expression for example is associated with lung, liver, and splenic macrophage identity, but is also associated with specific functions and metabolic states (Lavin et al., 2014; Varga et al., 2016). Most notably, several transcription factors implicated in macrophage polarization were enriched, including NR1H3 (liver X receptor alpha), NR4A2, ATF3, CEBPA, RUNX2, STAT1/2, and EGR2 (Bagnati et al.; Labzin et al., 2015; Lehtonen et al., 1997; Ramachandran et al., 2019; Satoh et al., 2013; Shaked et al., 2015; Veremeyko et al., 2018). While these transcription factors have been largely studied in isolation for their effect on macrophages (and other MNP types), enrichment of multiple factors suggests that dissecting higher-order interactions between these transcription factors may be fruitful.

**Figure 6.**
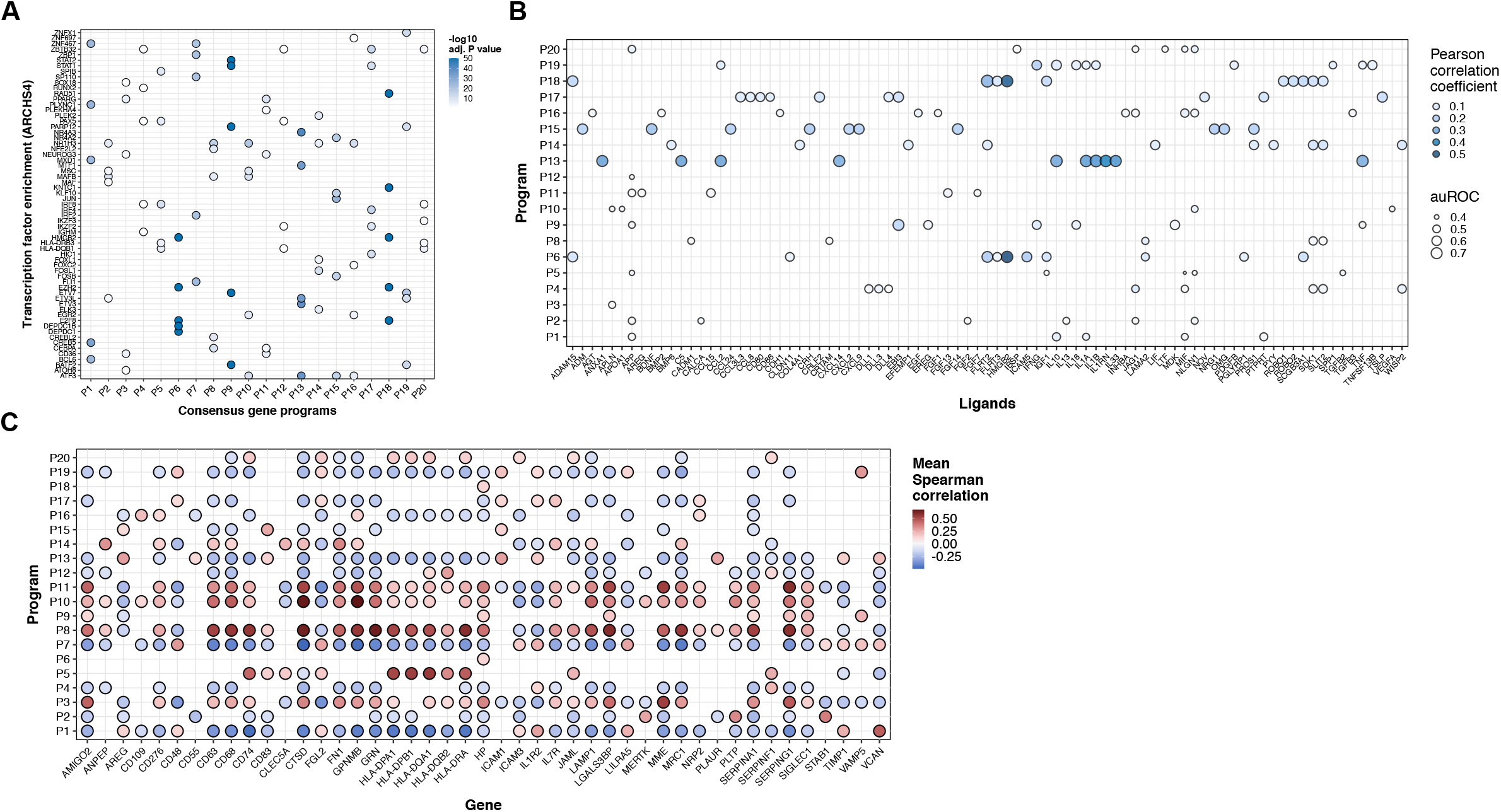
Enrichment of consensus gene sets suggests diverse molecular contributors and distinct roles. **(A)** Transcription factor coexpression enrichment for both program and marker gene sets using ARCHS4 identifies associated TFs with consensus GEPs. Top TF and corresponding adjusted P value is shown for each program. **(B)** NicheNet ligand activity analysis for programs identifies potential ligands driving program genes’ expression. Aggregated mean Pearson correlation coefficients represent ligand activity across all datasets for each program geneset. Only the top 2 MP clusters were used for each calculation. **(C)** Spearman correlation of annotated cell surface proteins with program scores averaged across datasets identifies associated surface markers. Cell surface proteins visualized are selected based on top 5 highest or lowest correlation per program.

To nominate potential ligand drivers of gene programs identified using NMF, we utilized NicheNet (Browaeys et al., 2020). For each dataset, we set the receiver population as the top 2 expressing clusters for each gene set and the sending populations as all other cells. These populations were used to define expressed target and background genes, enabling us to collate an average ligand signal (Pearson correlation coefficients) for each program gene set (Figure 6B) in each dataset, which was aggregated (**Methods**). This identified a range of known and potential ligands involved in MNP programming, such as ANXA1, IL10, IFNG, and IL33 (Ip et al., 2017; Lu et al., 2020; McArthur et al., 2020; Nathan et al., 1983). Interestingly, several hits included unappreciated ligands, such as JAG1, VEGFA, SPP1, EREG, and BMP2.

Lastly, cell surface proteins provide a means to isolate specific subpopulations for powerful, targeted analyses. To identify candidate receptors that differentiate cell states or indicate activity of gene expression programs, we utilized a reference of cell surface proteins to calculate the Spearman correlation of receptor counts with program scores (Figure 6C). This enabled an unbiased look at receptors that may be positively or negatively correlated with these broad descriptors of MNP heterogeneity. As expected, we identify frequent strong correlations with MHC class II receptors with various programs (P8, P10) and canonical MNP markers, such as CD63, CD68, CD83, MERTK. This analysis also reveals additional markers that have been briefly investigated, but are not typically considered. These markers, such as CD109, CD276 (B7-H3), CD48, and LILRA5, may be useful in describing MNP variation in future investigations (Alivernini et al., 2020; Lee et al., 2017; Rizzo et al., 2020; Ydens et al., 2020).

Overall, this collection of enrichment analyses provides an initial suggestion of which molecular factors could be responsible or associated with these reference cell states and programs.

## DISCUSSION

A wealth of single-cell RNA-sequencing atlases have enabled unbiased profiling of cellular populations across conditions in the lung. The myeloid compartment within these atlases has received varying annotation and no harmonization across these findings has been attempted. Moreover, while specific pathogenic expression signatures have been identified in individual studies, the application and validation of these signatures in other cohorts has yet to be conducted. We hypothesized that standardized processing and comparison of genes describing cluster-specific and cluster-agnostic variation could reveal highly conserved gene sets. Our analysis across 12 studies generated 17 marker and 20 program consensus gene sets. This reference offers a robust, fast, and interpretable framework to decipher the single-cell transcriptional states of mononuclear phagocytes within the lung.

We first took a cluster-based approach to defining consensus signatures. By splitting each published cohort into disease and healthy samples, we sought to correct for batch effects without over-correcting any variation potentially attributable to disease biology. After identifying granular clusters and retaining defining markers, the broad comparison reveals strong partitions representing similarly-defined populations. Although coexpression patterns and nuances may vary between communities, we find consistent marker genes across all communities defined within a partition. A complementary approach using the gene expression programs (defined from NMF factors) reveals similar robust partitions defined by consistent genes. Visualization of clusters or programs across datasets by these scaled metrics describe the unique features of these populations with no identifiable dataset effects. This offers the ability to project new programs or marker gene sets into this low-dimensional embedding of community-defining gene sets. Additionally, this approach avoids complex integration methods across datasets from different cohorts and combines information across individual cell profiles which can be noisy.

The combination of these complementary marker- and program-based approaches underscored concordant results, supporting these gene sets as dominant features of variation within the lung MNP populations in both healthy and diseased samples.

Careful inspection of these consensus gene sets revealed expected descriptions of major cell populations, such as alveolar macrophages or CD14+ monocytes, and rarer subsets, such as a fibrosis-associated gene set containing previously validated genes (Reyfman et al., 2019) and an inflammation-associated gene set related to poor cancer prognosis (Zilionis et al., 2019). These gene sets offer all the same utilities as the commonly used Gene Ontology resource, with the significant benefit of using well-detected and well-expressed genes captured using single cell technologies. We demonstrate these utilities by comparing gene set scores between healthy and disease samples, expanding upon the contribution of myeloid dysregulation to pulmonary diseases. M10 and P9 scores show an enrichment in disease samples in fibrosis and cancer, respectively, and should be utilized to further investigate the association of these genes and populations with disease. Moreover, identifying high P17-scoring cells enables the annotation of a rare immunoregulatory dendritic cell population recently described (Maier et al., 2020). While several interesting trends exist amongst all these data, these trends were of particular interest. These scores enable the easy interrogation of critical expression trends, analogous to fluorescence intensity shifts in flow cytometry analyses. These gene sets provide more context to individual genes typically focused on for validation in studies. Identifying how genes associate and behave similarly across contexts can provide new insight into gene function and gene-gene relationships in cell populations. Moreover, given the continuous nature of the MNPs broadly in single-cell RNA-sequencing data, correlation of these programs with an alternative measure of cell type similarity (the maximum likelihood calculated from a purified bulk reference) provides additional context to the interpretation of cells expressing these gene sets above baseline.

We further utilize this reference to dissect single-cell expression dysregulation in COVID-19 bronchoalveolar lavage samples. While two groups have described distinct clusters and proportional shifts in their analyses, we focus on expression shifts related to these gene sets, since they can be quickly interpreted and related back to all the studies analyzed in this study. Most notably, the M7 and P9 gene sets related to activated myeloid expression are dramatically increased in disease samples versus M5 and P11 related to resident macrophage expression increased in healthy samples. When we define a general threshold across all samples and calculate the percentage of cells above threshold, robust shifts in population structure, consistent between individual donors, is revealed. Overall, these findings provide complementary insights into the dysregulation of the myeloid compartment during COVID-19 infection. While we see a shift towards monocytes and monocyte-derived phagocytes based on expression patterns, we also see increased expression of distinct markers and programs found in other disease pathologies. This provides evidence for the dysregulation of mononuclear phagocytes in COVID-19 pathogenesis, specifically the shift away from resident, lipid-associated macrophages to inflammatory (defined by type 1 IFNs and inflammasome genes) monocyte-like cells.

While many opportunities still exist for interrogating these datasets broadly, as well as the gene sets defined here, we performed enrichment analyses to identify potential molecular factors. These analyses provided a host of molecules for follow-up including MNP-related TFs (e.g. NR1H3, CREB5, PPARG, SP110), active ligands (e.g. ANXA1, AREG, JAG1, IL1RN, SCGB3A1), and descriptive markers (e.g. LILRA5, MERTK, MRC1, PLAUR, SERPING1, SIGLEC1, STAB1). Cataloging *in vitro* behavior of these molecules could enable improved interpretation of these suggestive enrichment, NicheNet, and correlative results.

This study focused on identifying consistent trends across lung datasets that were robustly codescriptive of similar states and similar gene expression programs. We recognize that more subtle patterns in gene expression or coexpression may be lost and look forward to addressing this in future work. These profiles are limited by the sensitivity and type of technology used to generate these profiles, as well as the availability of raw count matrices and associated metadata. While we focused on the lung here due to the available data, scientific interest, and unmet need, understanding how these findings translate to other tissues remains unknown. Extending this work to other tissues and datasets is a priority. This raises the important question of how similar MNP populations are intra-niche versus inter-niche. Lastly, we recognize the enormous potential of many tools developed by the single cell community and expect that the application of these tools to these data can provide additional complementary and novel insights into the transcriptional landscape of these cells.

Collectively, these analyses provide a reference to interrogate past and future transcriptomics of monocytes, dendritic cells, and macrophages. While this study is focused on lung datasets, we imagine that these findings will be a useful starting point for studying the MNP compartment in other tissues. Robust and diligent compilation of cellular states from *in vivo* data provides the opportunity to more efficiently co-opt biological insights across disease contexts within a tissue. As additional attention is given to re-engineering innate immune cells therapeutically, this reference can serve to annotate states, identify new cellular and gene-gene relationships, and repurpose myeloid-directed therapies across diseases. We imagine a tight feedback loop between well-controlled *in vitro* and model *in vivo* modulation of these states with deeply defined human *in vivo* states to attempt to better define the disease and tissue signals responsible for these states. The continued generation and integration of single cell transcriptomics will provide a map to deciphering clinically-relevant *in vivo* cellular states.

## MATERIALS AND METHODS

### Resource Availability

#### Lead Contact

Further information and requests for resources and reagents should be directed to Bryan Bryson.

#### Materials Availability

This study did not generate new unique reagents.

### Data and Code Availability

Machine-readable files describing gene modules are available on Github (https://github.com/joshpeters/lungmps). Data used for these studies are available to download using the links in Supplementary Table 1. These datasets, annotated and formatted as Seurat objects (.rds), are available on Zenodo (https://doi.org/10.5281/zenodo.3894750).

### Experimental Model and Subject Details

This study did not use any new experimental models or subjects.

### Method Details

#### Clarification of Nomenclature

Given the confusing and variable usage of common nomenclature used in transcriptomic analysis, we define terms used in this work for clarity. Within each dataset, we define clusters, or groups of cells that have similar transcriptomic profiles in low-dimensional space. We describe these clusters with marker genes (“markers” for short) which are expressed and detected at higher levels in these clusters. Additionally, we define gene expression programs (GEPs, “programs” for short), which are the factors of matrix *W* describing the loadings of each gene generated from non-negative matrix factorization algorithms. These are sometimes referred to as “metagenes”. The generation and interpretation of these programs is described in previous work (Kotliar et al., 2019; Welch et al., 2019). Broadly, we refer to these markers or programs as gene sets. When we compare gene sets across datasets, we generate partitions, which represent larger groups of programs or markers defined from individual datasets. Ultimately, we describe these cellular populations and their transcriptional landscape by deriving both markers, derived from defining distinct clusters, and programs, derived from describing major axes of variation.

#### Preprocessing

Raw count matrices were downloaded from respective published sources. To ensure the gene names were harmonized as best as possible, they were mapped to the Ensembl database using the R package EnsDb.Hsapiens.v86. Gene names that did not map were searched using an HGNC reference of gene names, aliases, and Ensembl IDs. Identified Ensembl IDs were reconverted to gene symbols. Any unidentified gene symbols were left as is. Raw matrices were then split based on disease status for applicable datasets. Seurat objects were populated with these raw data and processed using the same procedure. First, cells were filtered based on technical covariates: < 20% mitochondrial reads, <50% ribosomal reads, <5% hemoglobin reads, <5% heat shock protein family reads, > 200 UMI counts, > 100 genes detected. Filtered cells were then batch-corrected (based on broad observation of batch effects across datasets) using Harmony (*RunHarmony*, Harmony R package) (Korsunsky et al., 2019).

#### Downstream dimensionality selection

20 downstream dimensions were used for each dataset. Several heuristics were calculated for each dataset, including identification of elbow point based on maximum distance to the line between the first and last components’ variance and the global maximum likelihood based on translated Poisson Mixture model using default parameters (*maxLikGlobalDimEst*, intrinsicDimension R package). Visual inspection in combination with these methods suggested between 10 and 20 dimensions for each dataset. For ease and consistency, we utilized 20 Harmony-corrected dimensions for downstream clustering for all datasets. As others have suggested, we reasoned that this accurate or slight over-estimation of dimensionality would work well universally across datasets.

#### Clustering and visualization

Cells were clustered using the Leiden algorithm using default parameters on the SNN graph utilizing the default parameters (*leiden_base_partition*, leidenbase R package). First, resolution values were scanned for parameters that generated between 5 and 30 clusters. 30 resolution parameters were then tested between this range to identify the value that generated the maximal modularity. This value is then used for an additional 30 iterations to determine a final clustering. Any clusters with less than 10 cells were merged utilizing a modified implementation of *GroupSingletons* (Seurat R package). To assess the potential number of clusters and agreement with Leiden results, cells were also clustered using the walktrap algorithm (*cluster_walktrap*, igraph R package). Clusters were renumbered based on the phylogenetic tree generated from averaged cluster expression and visualized using UMAP (*RunUMAP*, Seurat R package).

#### Global, general cell annotation

Cell clusters were annotated using a semi-supervised approach. First, we extracted high-confidence marker genes defined in a published, comprehensive healthy lung atlas (differentially-expressed genes, average log FC > log(2), BH-adjusted P value < 1E-10, top 20 based on average log FC) (Travaglini et al., 2019). Each label was expanded into cell class labels (e.g. immune, epithelial) and subclass labels (e.g. myeloid, lymphocyte) manually. Each cell was scored based on each set of DEGs and assigned a class, subclass, and cell type label based on its maximum score. Clusters were then annotated using a voting approach. We found this voting approach to be superior to a threshold-based approach using Gaussian mixture models. Cycling cells were annotated using a similar scoring approach with the cell cycle genes defined in Seurat. Lastly, cells were classified using the immunoStates reference using a Bayesian classifier described previously (Vallania et al., 2018; Zemmour et al., 2018; Zilionis et al., 2019). This classifier generates a probability for each cell and each type in the immunoStates reference. We summed the results from each macrophage label or dendritic cell label to define the overall macrophage or dendritic cell likelihood.

#### Mononuclear phagocyte gating and selection

Each dataset was initially classified using an automated approach, then reprocessed and cleaned with manual inspection of the measures described previously. First, we define a MNP score for each cell, which is the difference between the Travaglini scores for MNP cell types and non-MNP cell types. We then fit a Gaussian mixture model to this score and define an upper threshold of 2.5σ above μ. Clusters with a median score above this threshold were labeled as MP for the second round of classification. Cycling cells above this threshold were also included. Clusters that did not reach the median score threshold or contained more than 50% of other cell types (as classified by the maximum Travaglini score) were not MPs. This approach was repeated a second time on initial MP clusters. A second round of preprocessing and annotation revealed contaminating non-MP clusters that were manually inspected and removed. To aid in this procedure, a list of non-MP marker genes were defined for each dataset that included the highly predictive (auROC > 0.9) markers of non-MP clusters, lymphocyte receptor genes, and neutrophil genes (CSF3R, NAMPT). These excluded genes were used to identify contaminating clusters defined by excluded genes after reprocessing. These clusters are most likely due to ambient RNA distortion, suboptimal clustering, or unidentifiable heterotrophic doublets. While we cannot rule out the possibility that these cells are bona fide profiles representing a single cell expressing dominant and predictive transcripts of multiple clusters, further investigation is out of the scope for this study.

#### Unsupervised phagocyte classification

We utilized *scmap* (R package) to perform unsupervised projection of phagocyte clusters across all datasets, considering its speed and performance (Abdelaal et al., 2019; Kiselev et al., 2018) To compile a feature set for projection, we collated the marker genes for all clusters across all datasets and calculated their frequency and median auROC. We took the top 1000 genes that were identified in 4 or more datasets. These genes were then used to index each cluster from each dataset. The cluster indices (n = 290) were then projected to each dataset using a threshold of 0.3 to determine assignments. This procedure assigned a cluster label (or unassigned label) from each dataset for every cell within the query dataset. For each pair of query cells and reference dataset indices, we calculate the proportion of each reference assignment grouped by query clusters. This provides a voting-based similarity measurement for each pair of query and reference clusters. We sparsify these similarities by zeroing any proportions where the number of labeled cells is below 5 or the proportion is below 1%. We collate the similarities for each pair to generate a matrix of query clusters by reference clusters describing a complete bidirectional dictionary of cluster projections.

#### Generation of cluster-defining genes for marker comparisons

Filtered cells were clustered using the Leiden algorithm on the SNN graph, implemented in leidenbase (R package) utilizing the default parameters. First, resolution values were scanned for parameters that generated between 5 and 30 clusters. 30 resolution parameters were then tested between this range to identify the value that generated the maximal modularity. This value is then used for an additional 30 iterations to determine a final clustering. Any clusters with less than 5 cells were merged utilizing a modified implementation of *GroupSingletons* (Seurat R package). Clusters are then hierarchically clustered in the low-dimensional PCA space and renumbered for convenience. Mann-Whitney U and auROC testing is performed (*wilcoxauc*, Presto R package) and the second-to-max log fold change (LFC) is calculated, which is the fold change between the top two expressing clusters. Clusters that fail to be defined by gene markers (LFC > 1.25, second-to-max LFC > 1.05, AUC > 0.5, BH-adjusted P value < 1E-3) are then merged with the nearest cluster. Clusters are not merged if they are to be merged with multiple clusters (a node in the hierarchical tree) or a cluster defined by more than 50 markers. This prevented merging of poorly defined clusters with well-defined clusters, which could increase noise in the markers.

#### Comparison and clustering of gene sets

To directly compare gene sets that define clusters or factors, we utilize the Jaccard distance, as described previously (Kinker et al., 2019). We compute the symmetric Jaccard distance matrix and remove sets with no occurrences of a Jaccard similarity of more than 0.2. We perform agglomerative hierarchical clustering on this filtered Jaccard distance matrix and cut the tree to generate *k* partitions based on heuristic assessment of the within-sum-of-squares and silhouette plot across a range of potential *k* values. To assist in selection, we calculate a combined rank for each *k* by adding the sorted index of the within sum-of-squares and silhouette values and take the minimum as a suggestion for the minimal *k*. In practice, we set *k* on the higher end of potential values to generate more clusters and refine the partitions subsequently. For each partition, we then collate the original data (e.g. differential expression results, factor loadings) to generate consensus descriptions of each partition (see Methods sections **Generation of consensus markers**, **Generation of consensus gene expression programs**). We admiringly acknowledge that, during the course of this study, a method proposing a similar workflow with favorable results was published (Gao et al., 2019). This supports our approach and provides useful, additional bioinformatic tools for researchers.

Although attempted (not published), methods that integrate individual datasets are computationally prohibitive as cell population scales and are more difficult to interpret and validate, especially when looking at one cell type. By focusing on the easily interpretable Jaccard index and auROC metrics, we efficiently minimize study-specific effects and noise.

#### Generation of consensus markers

After generating partitions of cluster markers based on the Jaccard distances, the median AUC, median log FC, and number of contributing clusters for each gene was calculated. Resulting partitions were removed if they contained only 1 source dataset or were defined by gene families associated with technical covariates (see **Preprocessing**). Partitions consisting of one dataset were also removed after inspection. less We then filtered genes that did not appear in at least a third of contributing clusters (minimum of at least 2) and then sorted these by descending median AUC. The top 50 genes were chosen to represent an efficient, consensus gene set that defines clusters across datasets. This Jaccard-based approach was validated using additional marker gene sets and *scmap* (R package).

#### Integrative non-negative matrix factorization

Integrative non-negative matrix factorization was performed in LIGER as published(Welch et al., 2019). For each dataset, the MPs were integrated by batch across 30 dimensions with k = 20 and λ=5. K was chosen based on previous procedures (see Downstream dimensionality. This produced a usage matrix, H, a shared factor matrix, W, and donor-specific factor matrices V. This procedure was executed 30 times for each dataset. These replicates were compiled to generate robust W matrices using a modified procedure from consensus NMF(Kotliar et al., 2019). Briefly, all metagenes in all replicates (n = 600) were compiled into a single matrix and L2-normalized. The top 9 nearest neighbors were identified (ρ = 0.3). Factors beyond the 90th percentile of the average distance to the top 9 nearest neighbors were removed and the remaining factors were clustered on their loadings using K-Means (kmeans, stats) with 30 random starts to obtain 20 factor clusters. Grouped factors were compiled into a final set of factors by taking the median loadings. This consensus iNMF procedure generates a matrix W describing 20 donor-agnostic gene expression programs for each dataset used for downstream comparison.

#### Generation of consensus programs

Similar to generating consensus markers, factors were partitioned into groups using the procedure outlined previously (see **Comparison and clustering of gene sets**). Factors were defined by the top 5% of genes based on the non-zero loadings. The Jaccard distance was calculated and clustered using agglomerative hierarchical clustering with the Ward method (hclust R package, stats R package). Consensus genes were compiled based on their frequency in at least a third of the factors within each partition. Additionally, median consensus loadings were computed from respective factors within each partition for visualization.

#### Gene set scoring

We scored gene sets to generate a new aggregate expression level corrected for background expression of each cell, as described previously (Smillie et al., 2019; Stuart et al., 2019; Tirosh et al., 2016). In preprocessing, gene sets were scored using the *AddModuleScore* function in Seurat (R package). Downstream analyses utilized a modified version of this function in which scaled expression, not normalized expression, was used to aggregate the score. Briefly, gene sets were scored by aggregating the expression of each gene within the dataset minus the expression from control genes. For each gene, 30 control genes within the bin of expression (20 bins) for the gene of interest were selected.

#### Defining score thresholds and fraction of cells above threshold

After scoring cells for the broad expression of a gene set, we characterized the distribution of the scores amongst the cells of interest. We described the distribution of these scores as a bimodal mixture of Gaussian distributions (*normalmixEM*, mixtools R package) after visual and statistically inspection (diptest, Pearson diagram) where distributions were typically right skewed or bimodal. Given the difficulty of inspecting and fitting accurate models for every dataset and score, we utilized bimodal Gaussian distributions to capture cells either in the upper quantile of gene set expression or cells describing the second mode. We determine a threshold based on 2σ above the minimum peak μ in the fitted Gaussian mixture model. We also used the 90th quantile to determine shifts in scores. To do so, we bootstrapped the scores 1000 times from each condition label of interest and calculated the summary statistic distribution to report.

#### Comparison to M1 and M2 signatures

We derived M1-like and M2-like signatures from Martinez et al. and calculated their scores as previously described (Martinez et al., 2006). We report the Pearson correlation of these scores with program scores.

#### Enrichr enrichment analysis

We utilized Enrichr (enrichR, R package) to perform gene set enrichment analysis on our findings (Kuleshov et al., 2016). The ARCHS4, Reactome Pathway, and most recent GO databases were used (Ashburner et al., 2000; Lachmann et al., 2018).

#### NicheNet ligand activity

NicheNet was used to identify potential ligand-receptor activity within MNP populations, as outlined in the method vignettes (Browaeys et al., 2020). We define sender and receiver populations, background and target gene sets, and potential ligands. NicheNet returns ligand activities based on target genes relative to background genes. Potential receptors are then identified from top ligands. In this application, we defined sender populations as all non-MNP cells. The receiving population was defined as cells in the top two clusters based on the gene set score of interest. The Pearson correlation coefficient and auROC is reported as a measure of suggested ligand activity.

#### Surface protein-gene set correlation analysis

To identify surface protein expression correlated with gene set expression, we calculated the Spearman correlation of receptor counts with gene set scores. Putative surface proteins were selected based on the Cell Surface Protein Atlas (Bausch-Fluck et al., 2015).

## ACKNOWLEDGEMENTS

We thank the Blainey and Bryson laboratory members for valuable discussions and feedback, the researchers who produced the data used in this study and the donors who provided samples for these data, and the developers of the following R and Python packages: Seurat, Harmony, LIGER, cNMF, and the tidyverse collection.

J.M.P. and B.D.B were supported by grants R01A1022553 and R21DE026582. P.C.B. was supported by a Burroughs Wellcome Fund CASI Award.

## AUTHOR CONTRIBUTIONS

Conceptualization, J.M.P. and B.D.B.; Methodology, J.M.P.; Investigation, J.M.P.; Writing – Original Draft, J.M.P., B.D.B., and P.C.B.; Writing – Review & Editing, J.M.P., B.D.B., and P.C.B.; Supervision, B.D.B. and P.C.B.

## DECLARATION OF INTERESTS

J.M.P. and B.D.B. declare no competing interests. P.C.B. is a consultant to and/or holds equity in companies that develop or apply genomic, microfluidic, or single-cell technologies: 10X Genomics, General Automation Lab Technologies, Celsius Therapeutics, Next Gen Diagnostics, LLC, and Cache DNA.

## SUPPLEMENTARY INFORMATION

Supplementary Table 1. Study-level metadata for all scRNA-seq datasets used.

Supplementary Table 2. Sample-level metadata for scRNA-seq profiles used.

Supplementary Table 3. Consensus variable genes and associated metrics.

Supplementary Table 4. Pan-dataset cluster markers

Supplementary Table 5. Consensus gene markers from community partitions.

Supplementary Table 6. Pan-dataset programs

Supplementary Table 7. Consensus gene programs from factor partitions.

Supplementary Figure 1. Standard dataset processing metrics and results.

Supplementary Figure 2. Supporting analyses of consensus gene markers, related to Figure 2

Supplementary Figure 3. Supporting analyses of consensus gene programs, related to Figure 3

Supplementary Figure 4. Expression scores of consensus markers and programs in all clusters, related to Figure 4

---

**Figure S1.**
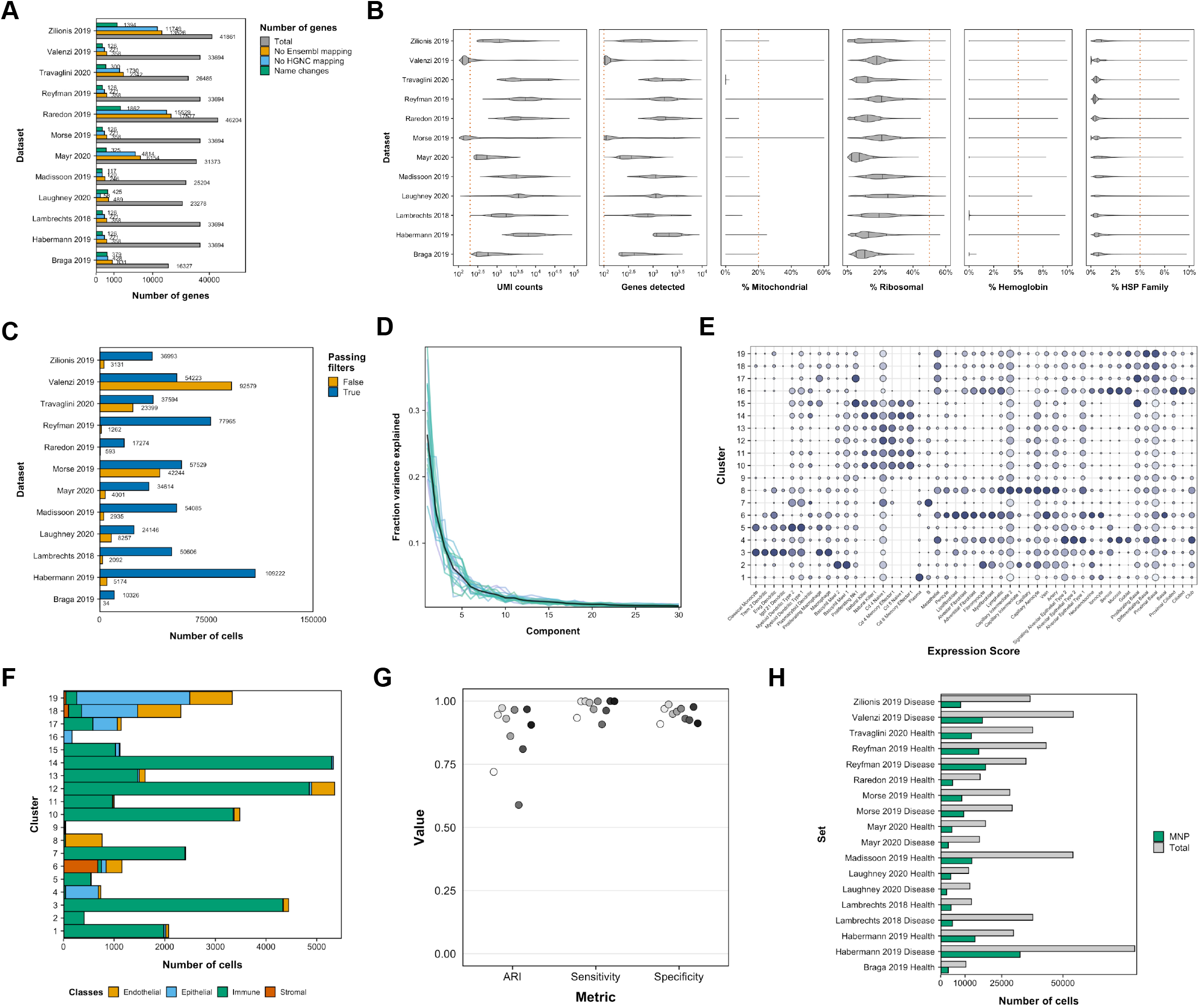
Datasets are processed uniformly to identify MNPs. **(A)** Number of genes mapping to current gene databases and changed. (B) Thresholds of each technical covariate for each study (n = 12). (C) Number of cells passing quality control filters from each dataset. (D) Fraction variance explained for all cells using Harmony-corrected components. Black line represents the average curve. (E) Example scores using differentially-expressed genes from Travaglini *et al.* 2020 applied to tumor samples from Lambrechts *et al.* 2018. (F) Cluster cell counts for each broad cell class from Lambrechts *et al.* 2018 tumor sample data. Each cluster comprises, to varying degrees of purity, cells classified as either endothelial, epithelial, immune, or stromal cells broadly. (G) Adjusted rand index, sensitivity, and specificity metrics for the agreement between classifying MNPs in this work versus published annotations where available. (H) Total number of cells analyzed and total number of MNPs identified from each dataset, split by disease status.

**Figure S2.**
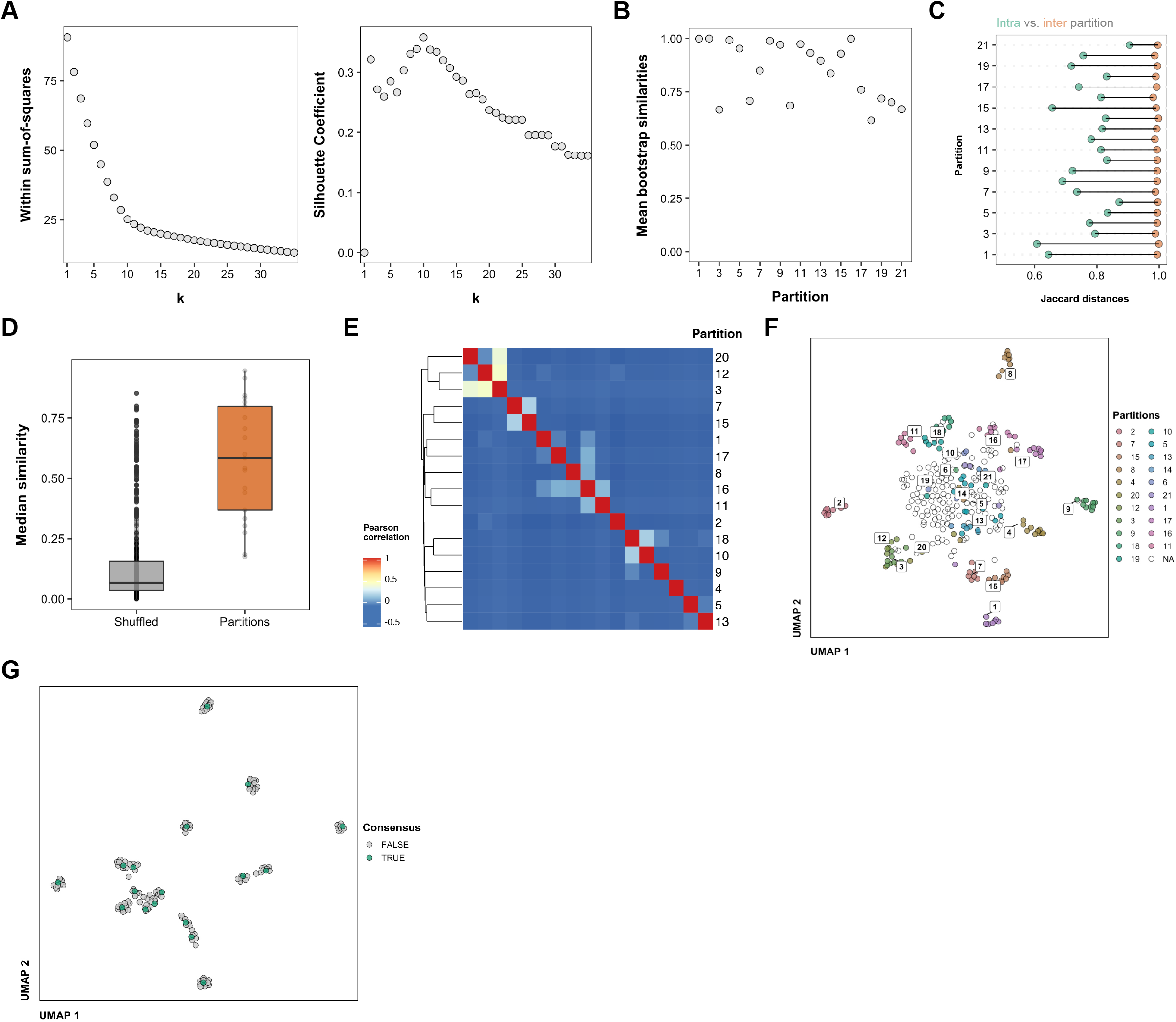
Consensus markers clustering metrics support distinct, robust partitions. **(A)** Within sum-of-squares distances and the silhouette coefficient for hierarchical clustering and resulting partitions (k = 21). **(B)** Mean bootstrap similarities across partitions (1000 replicates, clusterboot R package). **(C)** Average distance within partitions versus non-members for each partition. **(D)** Distribution of scmap similarities for clusters within partitions and random partitions sampled 30 times for each partition based on partition length. **(E)** Correlation of partitions based on median auROC values for defining genes. **(F)** UMAP embedding of all clusters from all datasets passing filters using auROC metrics for defining genes, colored by partition. NA clusters were not included in clustering based on below threshold similarity. **(G)** UMAP embedding of final clusters and final consensus markers. Consensus markers defined by median auROC values from each member cluster are shown in green.

**Figure S3.**
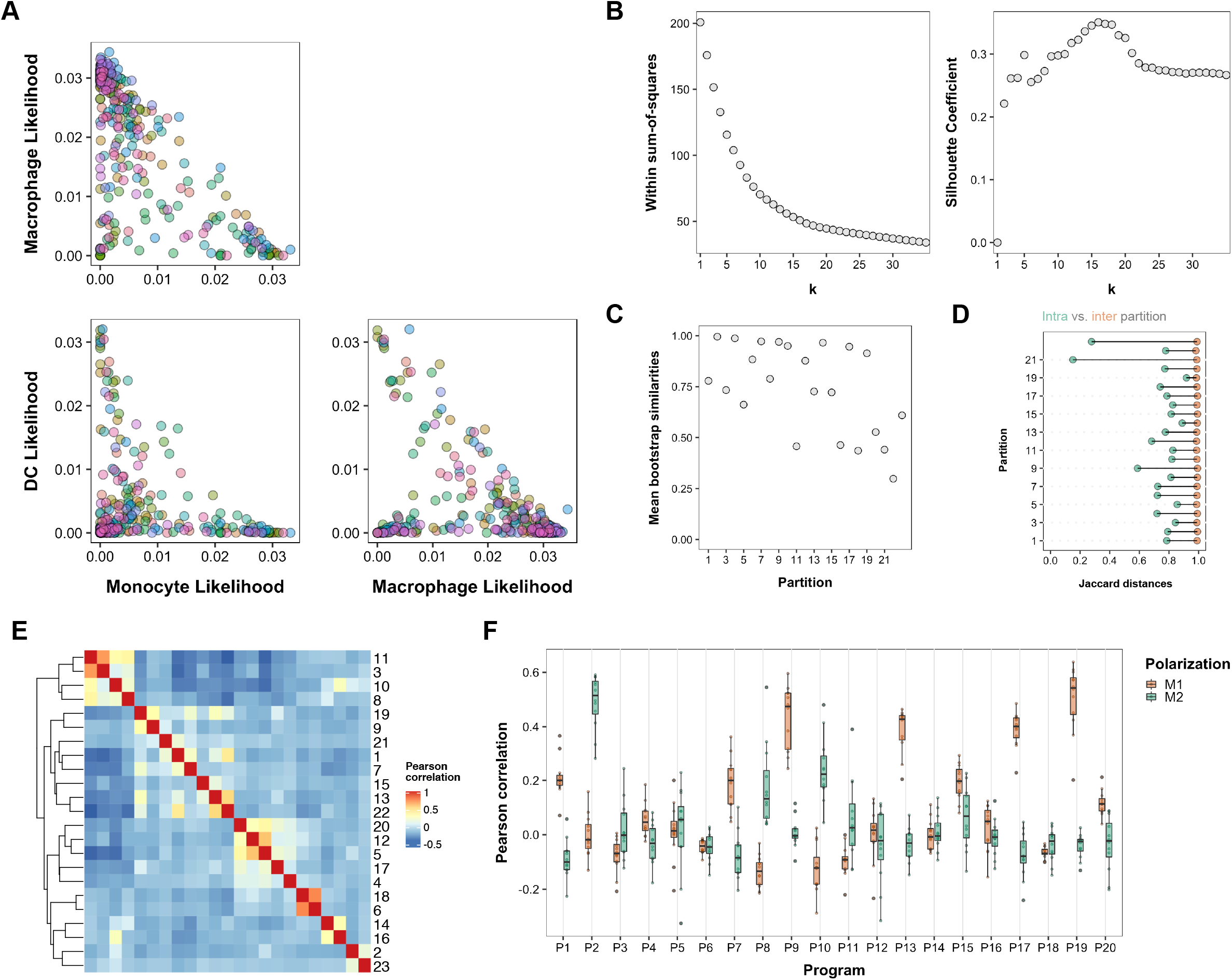
Consensus programs clustering metrics support distinct, robust partitions. **(A)** Monocyte, dendritic cell, and macrophage likelihoods plotted against each other for all clusters from all datasets, colored by dataset. Clusters fall along the ternary spectrum between each of these cell types. **(B)** Within sum-of-squares distances and the silhouette coefficient for hierarchical clustering and resulting partitions (k = 23). **(C)** Mean bootstrap similarities across partitions (1000 replicates, clusterboot R package). **(D)** Average distance within partitions vs. non-members for each partition. **(E)** Pearson correlation of partitions based on median auROC values for defining genes. **(F)** Pearson correlation of M1 and M2 gene signature scores with program scores. Each point represents one of the 6 studies used in Figure 4.

**Figure S4.**
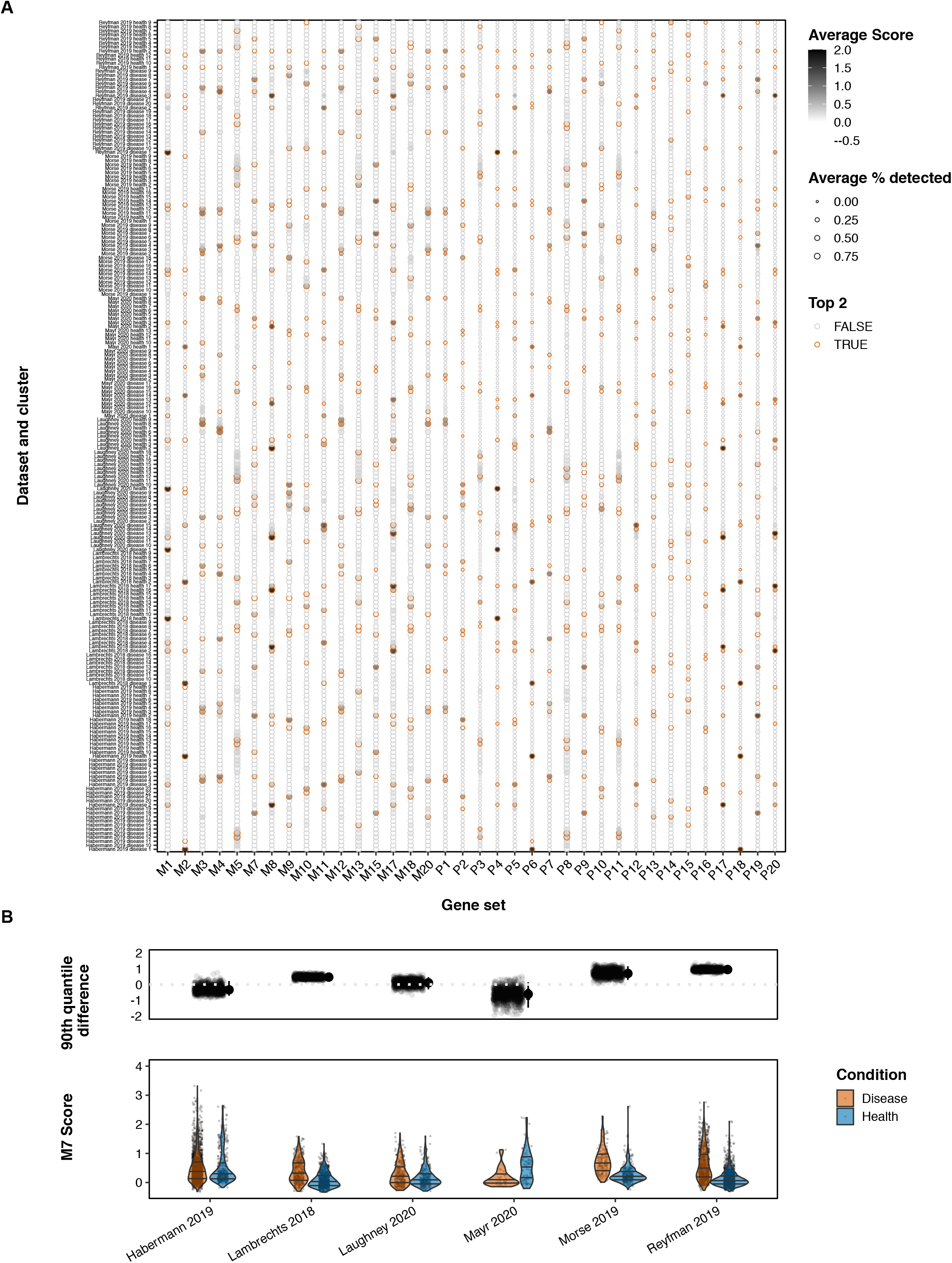
Consensus marker and program scores are variable across all clusters. **(A)** All marker and program scores for each cluster defined in the six comparative studies used in Figure 4. Darkness intensity represents higher expression of the gene set. The top two clusters for each gene set is highlighted with an orange outline. **(B)** M7 scores and bootstrapped 90th quantile differences for all six comparative studies.

